# Taxonomic scale dependency of Bergmann’s patterns: A cross-scale comparison of hawkmoths and birds along a tropical elevational gradient

**DOI:** 10.1101/2021.03.29.437440

**Authors:** Mansi Mungee, Rohan Pandit, Ramana Athreya

## Abstract

Bergmann’s rule predicts a larger body size for endothermic organisms in colder environments. The multiplicity of patterns and processes is expected because body size and temperature are two most fundamental factors on which many physiological, ecological and evolutionary processes depend, affecting all levels of biological organization, from individuals to communities. The confounding results from previous studies may be due to the differences in taxonomic (intraspecific, interspecific and community) and spatial (latitudinal vs elevational) scales. We compared Bergmann’s patterns for endotherms (Aves) and ectotherms (Lepidoptera: *Sphingidae*) along a same 2.6 km elevational transect in the eastern Himalayas. Using a large data spanning 3,302 hawkmoths (76 morpho-species) and 15,746 birds (245 species), we compared the patterns at the intraspecific (hawkmoths only), interspecific and community scales. At the interspecific scale, we account for phylogenetic non-independence in body mass by using a heirarchical linear mixed effects model for hawkmoths, and a phylogenetic generalised least squares model for birds. We assess the importance of using abundance-weighted metrics at the community scales, after accounting for spatial auto-correlation in communities. Hawkmoths exhibited positive Bergmann’s pattern at the intraspecific and abundance-weighted community scale. Intraspecific variation accounted for a substantial 33% variation at the community level. Contrary to this, birds exhibited a strong converse-Bergmann’s pattern at interspecific and community scales, both with- and without-abundance. Overall, all metrics which incorporate local traits and/or species abundances show stronger correlations than when this information is lacking. The multiplicity of patterns at a single location provides the opportunity to disentangle the relative contribution of individual- and species-level processes by integrating data across multiple nested taxonomic scales for the same taxa. We suggest that future studies of Bergmann’s patterns should explicitly address taxonomic- and spatial-scale dependency, with species relative abundance and intraspecific trait variation as essential ingredients especially at short elevational scales.

## Introduction

Bergmann’s rule is a popular, though contentious ecogeographical pattern which suggests a negative body size-temperature relationship in endothermic organisms. It is derived from a biophysical law which predicts better heat retention in larger-bodied animals due to their smaller surface area to volume ratio (Meiri & Dayan 2003). The direct and indirect influence of body size in many physiological, macroecological, and evolutionary processes (Bartholemew et al. 1981; Gillooly et al. 2005) and the simplicity of the initial explanation for this relationship are perhaps responsible for its popularity in ecological studies (Meiri & Dayan 2003; Watt et al. 2010; Olalla-Tárraga 2011). However, the results till date have been mixed, spanning positive, negative, and no correlation (Ashton 2002; Blackburn & Hawkins, 2004; Rodríguez et al. 2008; Olson et al. 2009; Shelomi 2012; Gutiérrez-Pinto et al. 2014; Gohli & Voje 2016; Freeman 2017; Nwaogu et al. 2018; Riemer et al. 2018).

In ectoherms, a converse Bergmann’s pattern has been proposed due to the shorter seasons associated with higher latitudes and elevations, which limits developmental time and hence body size (Atkinson & Sibly, 1997; Mousseau, 1997; Chown & Gaston, 2010). However, even here, a variety of results have led to the development of several taxon- and life history-specific mechanisms including heat conservation (Zamara-Camacho et al. 2014). Some of the commonly hypothesised mechanisms, in both ectotherms and endotherms, include the starvation resistance hypothesis (Cushman et al. 1993), growing season length (Bidau & Marti 2007), water availability (Ashton 2002), converse water availability (Zug et al. 2001), primary productivity (Rosenzweig 1968), energy-water conservation (Olalla-Tárraga et al. 2009), and competition and prey-predator dynamics (McNab 1971).

The primary focus of research on the body size-temperature relationship has been to assess the relative importance of these different processes across taxa and regions. However establishing the generality of a process, even for the same organismal group, has been hindered by the differences in the taxonomic scale (intraspecific, interspecific or community) at which studies are conducted (Gaston et al. 2008, Olalla-Tárraga et al. 2010; Figure 1). Studies at the intraspecific scale, also known as the population-approach, have documented clines along a temperature gradient (Ashton 2002; Yom-Tov & Geffen 2006; Fan et al. 2019) whereas interspecific studies have examined the pattern across multiple species using mean values for both body size and temperature (e.g. Freeman 2017; Alhajeri & Steppan 2016; Beck et al. 2016). The less common community- or assemblage-approach uses mean body size across all co-occurring species observed within a pre-defined area such as an elevational belt or a latitudinal grid (e.g. Blackburn & Gaston 1996; Olalla-Tárraga et al. 2006; Rodríguez et al. 2008; Zeuss et al. 2017; Brehm et al. 2019).

**Figure 1.**
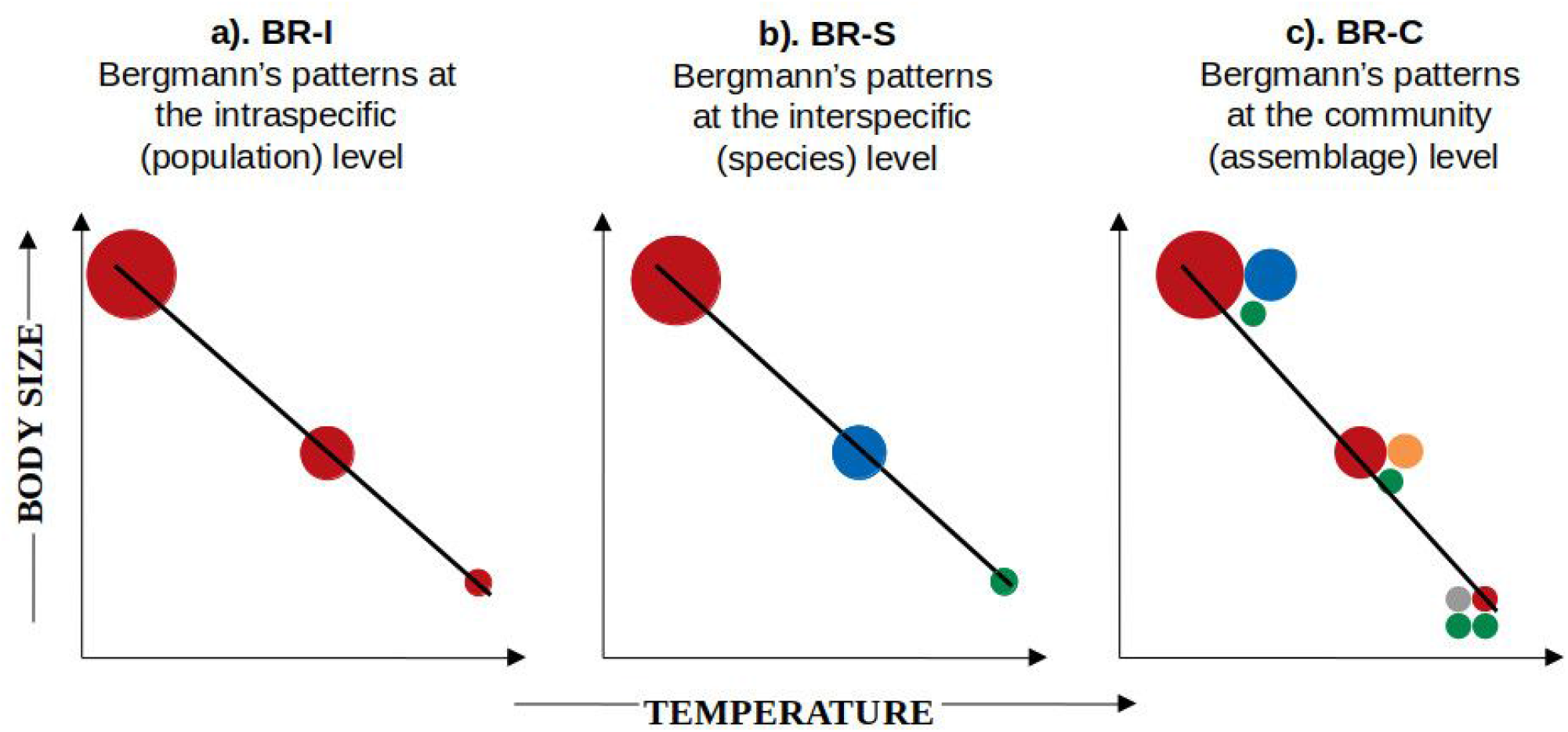
Three hypothetical scenarios are shown to illustrate a positive Bergmann’s pattern at three xonomic scales: intraspecific (BR-I), interspecific (BR-S) and community (BR-C). Each circle presents an individual, colors represent different species, and the size of the circle is proportional to e body size of the individual. a). **BR-I:** In the intraspecific approach, individuals of the same species re measured at different points along a temperature gradient, and individuals observed at higher mperatures exhibit lower body size; **b). BR-S:** In the interspecific approach, multiple species are corded along a temperature gradient, and overall smaller species are observed at higher temperatures; **b). BR-C:** In the community approach, all individuals within a pre-defined grid (or latitudinal / evational belt) are measured and community mean body size is lower at higher temperature. The BR-pattern can occur due to i). *Intraspecific variation* (e.g. individuals of red species are becoming maller with increasing temperature); ii). *Interspecific variation* (e.g. blue species is replaced by the maller orange species, which is further replaced by the even smaller grey species), or due to iii). *ariation in species abundances* (e.g. the green species of constant body size is found across the breadth f the gradient, however its abundance is higher in the community at higher temperatures).

Studies of the intraspecific pattern usually focus on a single or few closely-related species which exhibit large environmental (temperature) ranges (Ashton 2002; Gutiérrez-Pinto et al., 2014; Freeman 2017). Thus, although critical for understanding species-specific responses, the intraspecific approach may not be ideal for addressing the generality of an ‘eco-geographical rule’ since a majority of species do not span a sufficient range of temperature to exhibit marked intraspecific variation in body size (Gaston et al. 2008). The interspecific approach, on the other hand, uses information on multiple species, but constraints the data to species mean body size and mid-point of its elevational/latitudinal distribution. This ignores intraspecific variation, relative species abundances and range overlap between species - all three of which are shown to be strong determinants of macroecological patterns (Blackburn & Hawkins 2004).

Several authors have highlighted the scale-dependency of Bergmann’s patterns, and the community-approach is often remarked as the most appropriate way to investigate body size variation along broad temperature gradients as it can account for variations at both intra- and interspecific levels (Blackburn & Hawkins 2004, Meiri & Thomas 2007; Olalla-Tárraga et al., 2007). In the context of a Bergmann’s pattern, a greater contribution of intraspecific variation or change in species abundances to the variation in community mean body size may be indicative of a stronger role of size-related competition in facilitating species coexistence within a community (MacArthur & Levins 1967; Siefert et al. 2015). On the other hand, stronger interspecific variation, i.e. species turnover, is associated with strong environmental filtering on species’ body-, and therefore range-size (Keddy 1992; de la Riva et al., 2015). More importantly, the variations at the intra- and interspecific scales may also occur in opposite directions, resulting in a mis-characterization of the pattern or the process when either information is lacking (Vellend et al. 2014). Indeed many studies in plant functional ecology have shown a congruence of patterns and processes across different taxa only when individual-level data were taken into consideration (Kichenin et al. 2013; Siefert et al. 2015; Des Roches et al. 2018). Despite this, we found only two community level investigations of Bergmann’s patterns which have incorporated both, intraspecific variation and species relative abundances – one on bees (Classen et al., 2017) and another on moths (Brehm et al., 2019).

Body size is arguably the most important animal trait, and therefore it is not surprising that its variation is determined by several different processes, usually in a case- and taxon-specific manner. However, more importantly, the observed “diversity of Bergmann’s patterns” and the associated processes is also influenced by the different contexts of the many studies. They have differed not only in their taxonomic (intraspecific, interspecific or community) and spatial (latitudinal or elevational) scales, but also in the metrics used for the same scales (e.g. with and without species abundances/intraspecific variation for the community scale along elevational gradients; Beck et al. 2016; Brehm et al. 2019). Here, we investigate whether reducing the number of confounding factors would make for greater uniformity of body size-temperature relationships than seen in previous cross-taxa and cross-scale comparisons. We compare body size variation of endotherms (Aves) and ectotherms (Lepidoptera: *Sphingidae*) along the same compact elevational transect spanning 200-2770 m in the eastern Himalayas. We hypothesize that if heat conservation is the primary determinant of variation in body size, then we should observe a positive Bergmann’s pattern in both, hawkmoths and birds, and at all three taxonomic scales – intraspecific (only hawkmoths), interspecific and community, i.e. body size should increase with elevation at all taxonomic scales and for both organismal groups. We further assess the importance of species relative abundances and/or intraspecific variation in determining community level variation in body mass for the two taxa.

Sphingidae, commonly known as hawkmoths, are relatively easy to identify to the species level, and comprise over 1600 species globally with the highest diversity in the tropics (Beck et al. 2006a; 2006b, 2009; Kitching 2020). Birds are popular targets of studies of Bergmann’s pattern due to their high and easily identifiable diversity, well documented geographical distributions (Orme et al. 2006; Quintero & Jetz 2018), morpho-trait measurements (Ali & Ripley 1972; Rasmussen & Anderton 2005; Wilman et al. 2014), evolutionary relationships (Jetz et al. 2012), and perhaps their role in the original study (Salewski & Watt 2017).

## Materials and Methods

### Study region and the data

The study area and the hawkmoth data are described in Mungee & Athreya (2020a; 2020b). Briefly, hawkmoth individuals were sampled at 13 elevations separated by approximately 200 m between 200 and 2770 m in Eaglenest Wildlife Sanctuary in the eastern Himalayas of Arunachal Pradesh, northeast India. The region is one of the world’s 8 ‘*hottest hotspots*’ of biodiversity and endemism (Myers et al., 2000) with a rich avifauna (Orme et al. 2005; Price 2012). A novel photogrammetric method was used for morphological measurements of individuals of (unfettered) hawkmoths which settled on a UV light screen (Mungee & Athreya 2020a).

We carried out transect counts of birds along the same elevational gradient, but with a higher elevational resolution of 50 m. Each elevational transect was sampled on 12×2=24 occasions during 0600-1200 hrs in April-June, 2012-2014. We minimised systematic differences in bird activity across the 6-hr sampling window by subdividing it into three periods of 2 hour each – early (0600-0800 hrs), mid (0800-1000 hrs) and late (1000-1200 hrs). The 12 replicates at each elevation were equally distributed across these 3 periods. We could not sample below 500 m inside Eaglenest due to accessibility issues. Therefore, we sampled the lowest elevation (i.e. 200 m) in the adjoining Pakke Tiger Reserve, about 20 km from the 500 m transect in Eaglenest. Species mean body mass were obtained from published resources (Ali & Ripley 1972; Rasmussen & Anderton 2005; Schumm et al. 2020). Due to a strong linear relationship between temperature and elevation across our steep study site (r^2^ = 0.98; p < 0.001), we use elevation as the independent variable in all analyses (Mungee & Athreya 2020b).

### Statistical analyses

#### Intraspecific Bergmann’s pattern (BR-I)

We investigated the intraspecific pattern only for hawkmoths since we did not measure traits of individual birds. We estimated the pattern for hawkmoths as a group by using normalised body mass and elevation with species mean values:

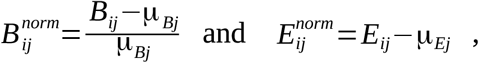

where B is body mass, E is elevation, µ is the species mean, and the indices indicate the *i-th* individual of the *j-th* species. With this normalisation we were able to pool the data from all individuals regardless of species and elevation to derive the intraspecific relationship between body mass and elevation for the ‘community’ as a whole. We also examined the intraspecific pattern for individual species which had more than 20 individuals and spanned more than 800 m (sampled at 4 or more elevations) in elevational range.

#### Interspecific Bergmann’s pattern (BR-S)

For the interspecific analysis, the species’ mean body mass (mean across all individuals from all elevations for hawkmoths; published mean values for bird species) was determined as a function of the midpoint of the elevational range for the species using ordinary least squares regression. Following previous practice we used log_10_(body mass) as the dependent variable at this level. However, body mass may be phylogenetically conserved across species leading to non-independence of residuals in a linear regression. To account for this, we used different approaches for hawkmoths and birds depending on the availability of information on phylogenetic relatedness among species.

In case of hawkmoths, we used the phylogenetically informed taxonomic classification assembled by Beerli et al. (2019) to perform a stepwise hierarchical linear mixed effect model with subfamily, tribe and genus as random effects and elevation as the fixed effect. Response (body mass) and explanatory (elevation) variables were standardized to zero mean and unit variance to make the coefficient estimates comparable within and between the hierarchical models. In case of birds, we used the phylogenetic tree provided by Schumm et al. (2020) for birds of eastern Himalayas. We added 13 missing species to this tree using the latest global avian phylogeny (Jetz et al. 2012). To do this, we scaled their branches to convert them into ultrametric trees, and used the relative branch lengths and placement from the global phylogenetic tree. We ran a phylogenetic generalized least square (PGLS) regression and estimated the phylogenetic signal of body mass by maximum-likelihood estimated values of Pagel’s lambda (Pagel, 1999).

#### Community-scale Bergmann’s pattern (BR-C)

For the community approach (BR-C), we calculated three metrics to assess the importance of species abundances and/or intraspecific variation in characterizing the community-level variation in body mass of hawkmoths and birds:

i. *CM-S (Community mean using species data):* Mean body mass across all co-occurring species at an elevation without considering their relative abundance. For hawkmoths, a single ‘regional’ mean (across all individuals from all elevations) body mass value was used for each species; this is the usual method while assessing grid-level assemblage patterns across latitudinal gradients (Olalla-Tárraga et al., 2010); it depends only on turnover in species richness *i*.*e*., species occurrences.
ii. *CM-A (Species Community mean with species abundance):* Mean of species mean trait value, weighted by elevation-specific abundance. This has been usually termed Community Weighted Mean (CWM) in previous studies (Cornwell & Ackerly 2009) but we suggest that the terminology used here brings out the differences in the three metrics in an easier manner; this metric encapsulates turnover in species diversity i.e., both species richness and abundance.
iii. *CM-I (Community mean using individual data; only hawkmoths):* Mean of all co-occurring individuals at an elevation, regardless of their species; it includes turnover in richness and abundance, and intraspecific variation.

We calculated the linear regression of the BR-C metrics with elevation, with and without weighting the communities using species richness (following Meiri & Thomas 2007). Additionally, for the metric CM-A, we also compare the patterns using community-weighted median, instead of mean. For all community level plots, we evaluated spatial autocorrelation in mean body mass by using Akaike Information Criterion (AIC) and the Likelihood Ratio (LR) tests of residuals (Diniz-Filho et al. 2003).

For hawkmoths, we also assessed the relative contributions of intra- and interspecific variability effects on community mean body mass along the elevational gradient following the variance decomposition method proposed by Lepš et al. (2011). This method is based on the decomposition of the total sum of squares (SS_specific_) of the community-level trait variance along an environmental gradient (elevation) into three additive components: ‘fixed’ (SS_fixed_), ‘intraspecific’ (SS_intraspecific_) and ‘covariation’ (SS_cov_) effects:

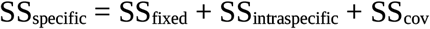

The two community-mean metrics used for this decomposition i.e. ‘fixed’ and ‘specific’ averages correspond to our CM-A and CM-I metrics. The ‘specific’ community-average body mass is calculated using the ‘local’ species body mass i.e. mean across individuals measured at that elevation (thus includes both inter- and intraspecific effects), and ‘fixed’ community-average body mass uses ‘regional’ species body mass i.e. mean across all individuals from all elevations along the gradient (includes only the interspecific effects). The effect of ‘intraspecific’ community mean is then calculated as the difference between ‘specific’ and ‘fixed’ averages. Linear regressions of the ‘specific’, ‘fixed’ and ‘intraspecific’ community averages are assessed, with elevation as an explanatory variable. The sums of squares for each of the three community-average measures explained by elevation is then used to assess the relative contributions of the intra- and interspecific components (Lepš et al., 2011). SS_cov_ component reflects the effect of covariation between inter- and intraspecific trait variability. We used 999 bootstraps to assess the significance of the three components.

All statistical analyses were performed in R 3.6.3 (R Core Development Team 2013) and the following packages were used: *spdep 1*.*1*.*3* (Bivand et al. 2018), *vegan 2*.*5*.*6 (*Oksanen et al. 2019), *cati 0*.*99*.*3* (Taudiere & Violle 2015), *ape 5*.*3* (Paradis & Schlip 2018), *lmerTest 3*.*1*.*3* (Kuznetsova et al. 2017), *caret 6*.*0*.*86* (Kuhn 2020).

## Results

We recorded a total of 4,731 hawkmoth individuals spanning 80 morphospecies, 30 genera and 3 subfamilies (Figure S1; Table S1). We measured traits for 3,302 individuals from 76 species; the remaining either did not sit on the gridded screen or could not be measured due to poor image quality. We have presented the patterns at all taxonomic scales using only the “trait” sample of 3,302 individuals, since we wished to compare results across taxonomic levels with the same data. Adding the rest of the sample, i.e. abundance-weights for the CM-A metric in the community approach, from the 4,731 individuals data did not change the results in any significant manner (Supporting Information Figure S2). We recorded 15,746 individual birds from 245 species, 150 genera and 50 families (Table S1).

### BR-I

At the pooled intraspecific level, hawkmoth body mass showed a weak, but significant positive correlation with elevation (Table 1; Figure 2a; r^*2*^ *= 0*.*001, p < 0*.*05*). The intraspecific patterns of the 24 species in our data which had an elevational range of 800 m or more and had more than 20 individuals were idiosyncratic (Table S2; Figure S1). Of them, the patterns of 17 (71%) species were not statistically significant, and only 6 (25%) species (*Acosmeryx naga, Acosmeryx omissa, Cechetra lineosa, Callambulyx rubricosa, Eupanacra sinuata* and *Rhagastis lunata)* exhibited weak but positive correlations, and one species a negative correlation (*Theretra clotho*) (Table S2; Figure S1).

**Table 1.**
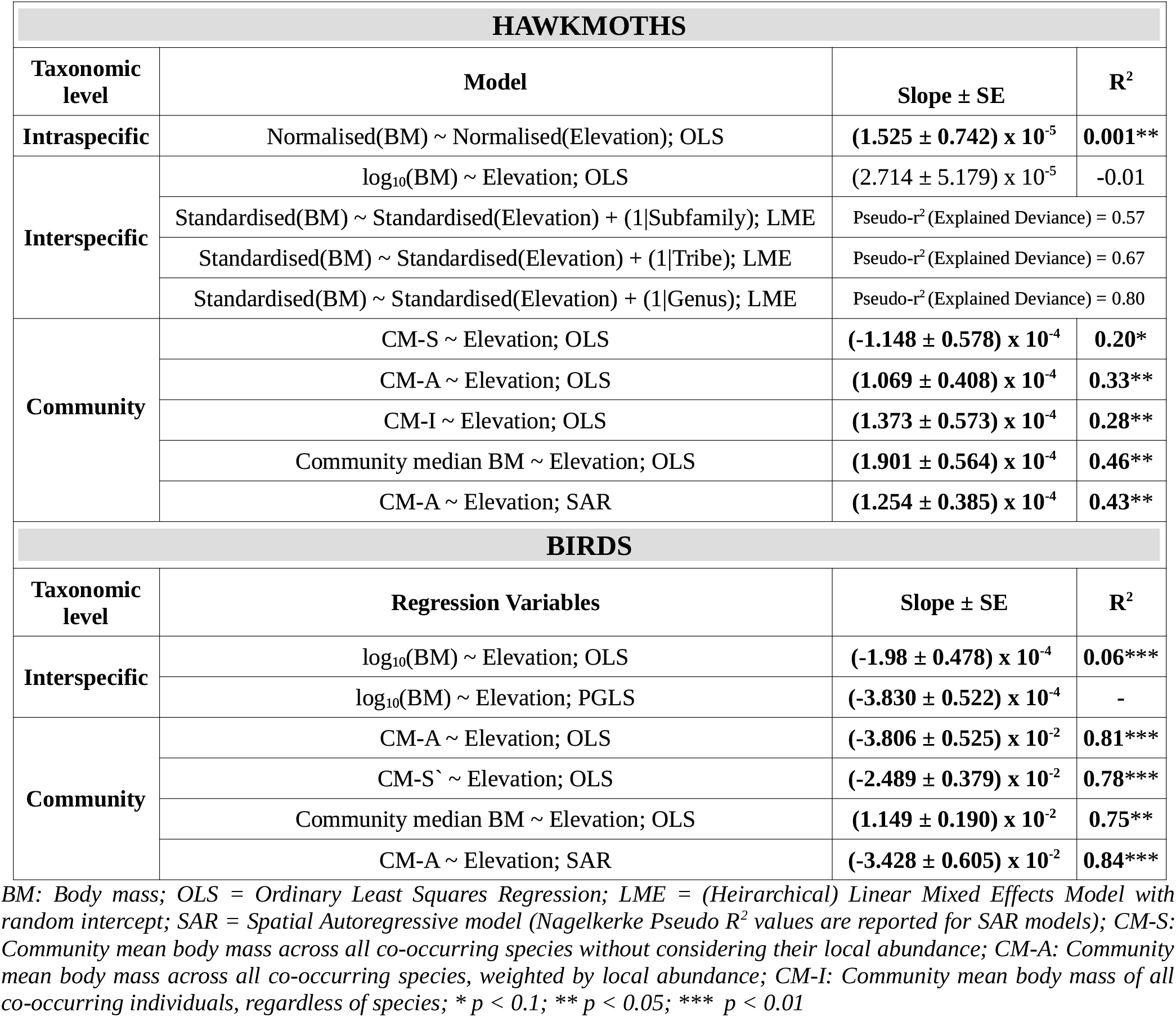
Linear regression between body size variation and elevation for hawkmoths and birds, at ifferent taxonomic levels. All slopes significant at p < 0.1 are in bold.

**Figure 2.**
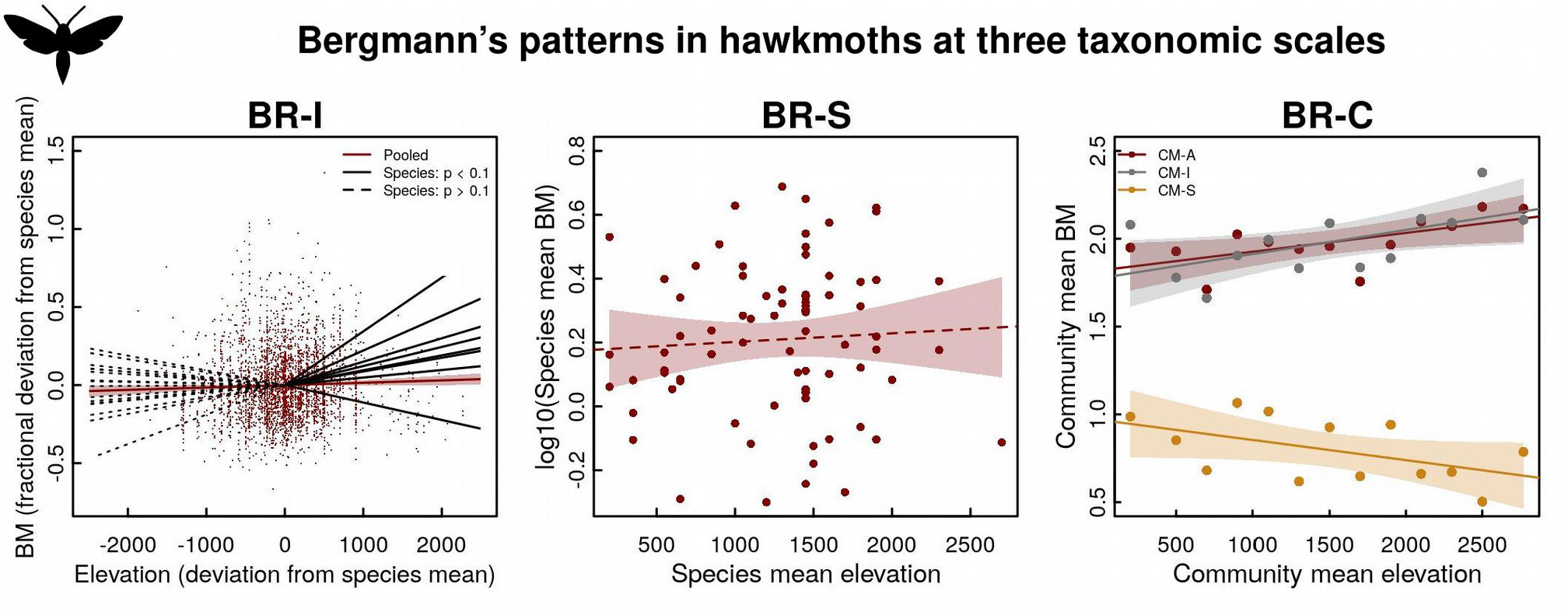
Relationship between elevation and body mass (BM) for hawkmoths at different taxonomic cales. ***a) BR-I:*** average intraspecific pattern obtained by pooling data of all individuals from all pecies. The red line represents the regression fit for the pooled data, while the black lines represent the 4 individual species which were sufficiently numerous (also see Figure S1); ***b) BR-S:*** interspecific attern using species mean body size and midpoint of the elevational range; ***c) BR-C: (i) CM-S*** *Community mean using species data):* Mean body mass across all co-occurring species at an elevation ithout considering their relative abundance (orange), ***(ii) CM-A*** *(Species Community mean with pecies abundance):* Mean of species mean trait value, weighted by elevation-specific abundance (red); (***iii) CM-I*** *(Community mean using individual data):* Mean of all co-occurring individuals at an evation, regardless of their species (grey). Regression coefficients for all the linear fits are listed in able 1. All fits significant at p < 0.1 are shown as solid lines, the rest are as dashed lines.

### BR-S

We found no support for an interspecific Bergmann’s pattern in hawkmoths using ordinary least squares regression (Table 1; Figure 2b; r^*2*^ *= -0*.*01, p = 0*.*6)*. Across the three hierarchical models, the explained variance (*pseudo-r*^*2*^) was consistently high (>0.5) and increased with each nested level (*subfamily = 0*.*57; tribe = 0*.*67; and genus = 0*.*80*). Overall, integration of the phylogenetic information did not affect the significance of the BR-S pattern (*p = 0*.*12 – 0*.*67*). Contrary to hawkmoths, we found a significant converse-Bergmann’s pattern for birds at the interspecific level (Table1; Figure 3a; *r*^*2*^ *= 0*.*06, p < 0*.*01*). Phylogenetic generalised least square (PGLS) models gave similar results (Table1; *λ = 0*.*88;* 95% confidence interval = [0.85, 0.90]), hence we discuss only OLS results below.

**Figure 3.**
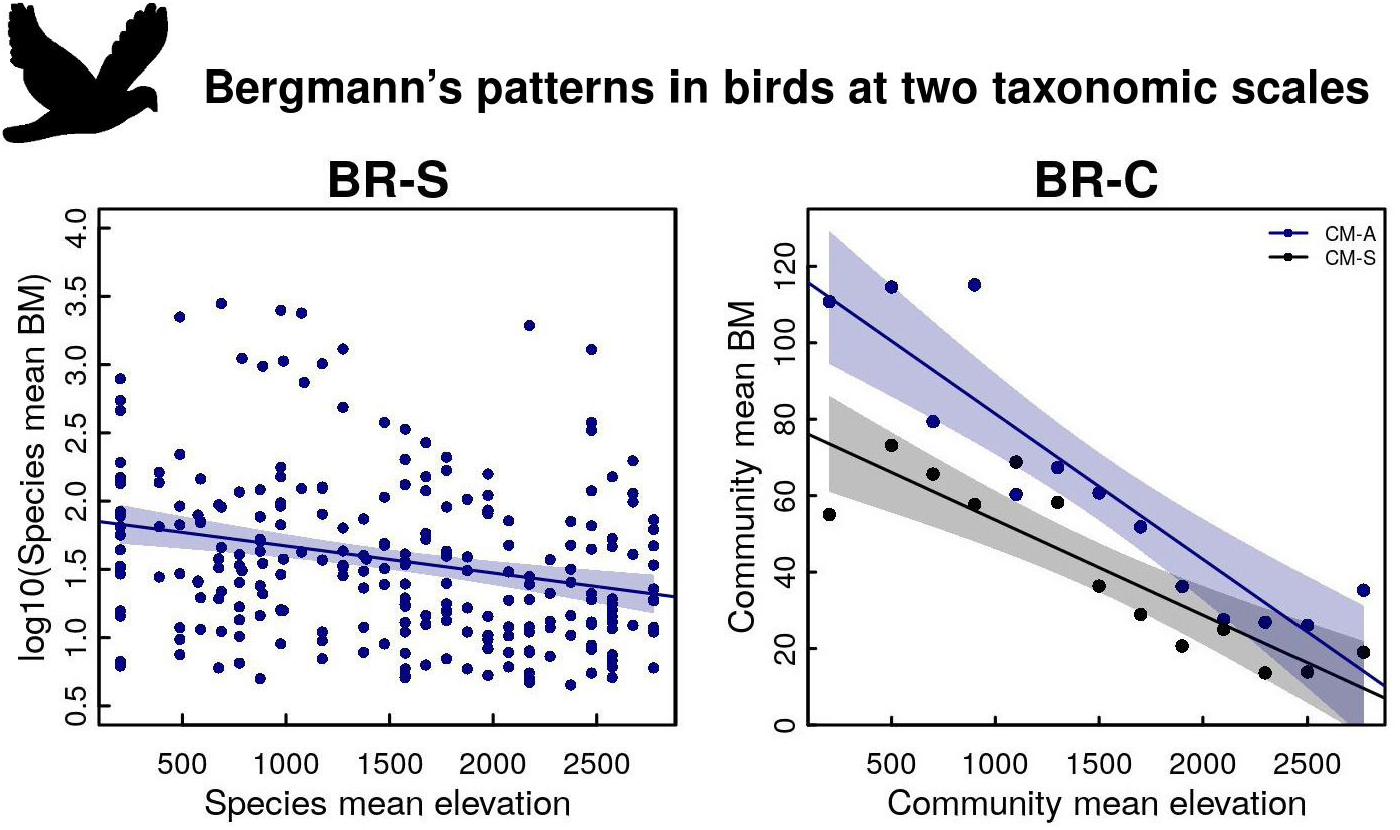
Relationship between elevation and body mass (BM) for birds at two taxonomic scales. ***a) R-S:*** interspecific pattern using species mean body size and midpoint of elevational range; ***b) BR-C:*** *(i) CM-S (Community mean using species data):* Mean body mass across all co-occurring species at an evation without considering their relative abundance, *(ii) CM-A (Species Community mean with pecies abundance):* Mean of species mean trait value, weighted by elevation-specific abundance; gression coefficients for all the linear fits are listed in Table 1. The regression lines were all significant p < 0.1.

### BR-C

We found a significant positive Bergmann’s pattern in hawkmoths at the community level for the abundance-weighted metric without incorporating intraspecific variation i.e. CM-A (Table 1; Figure 2c; *r*^*2*^ *= 0*.*33; p < 0*.*05*) and for the abundance-weighted metric which incorporates intraspecific variation i.e. CM-I (r^*2*^ *= 0*.*28; p < 0*.*05*). However, surprisingly, we found a significant negative pattern using the metric CM-S which does not incorporate ‘local’ abundance or intraspecific variation (Table 1; Figure 2c; r^*2*^ *= 0*.*20; p < 0*.*1*). Similar to the interspecific analysis, community mean body mass exhibited a significant converse-Bergmann’s patterns in birds (Figure 3b). Unlike in hawkmoths, the pattern was consistent between CM-A (*r*^*2*^ *= 0*.*81; p < 0*.*01*) and CM-S (*r*^*2*^ *= 0*.*78; p < 0*.*01*). Weighting the communities by species richness (Table S3), or using median instead of mean (Table 1) did not significantly affect the parameter estimates, for both hawkmoths and birds. Variance partitioning of hawkmoth community mean body mass showed that interspecific variation (STV) and intraspecific variation (ITV) contributed 54.2 ± 8.1% and 33.0 ± 5.5% respectively, with an overall 12.8 ± 8.4% positive co-variation between the two (Figure 4).

**Figure 4.**
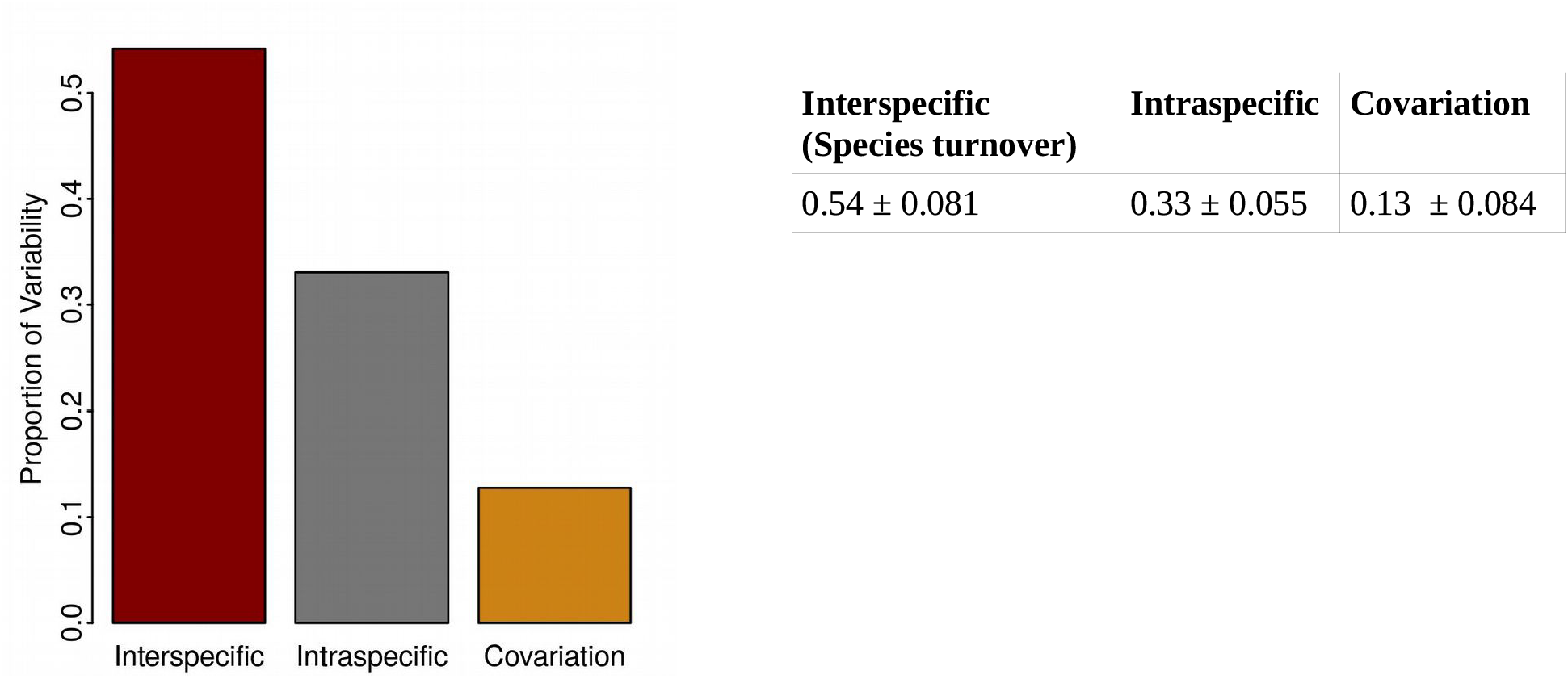
For hawkmoths, the relative contribution of intraspecific trait variation, species turnover (richness + abundance), and their covariation was assessed using the variance partitioning approach proposed by Léps et al. 2011.

## Discussion

We have shown that hawkmoths and birds exhibit contrasting body size-elevation relationships at the abundance-weighted community level along the same 2600 m elevational transect in the east Himalayas. Interestingly, endothermic birds exhibited a strong converse-Bergmann’s pattern while ectothermic hawkmoths showed a positive Bergmann’s pattern. Statistically, the patterns varied across different taxonomic scales within each group, although qualitatively they were similar only in case of birds. Hawkmoths exhibited significant Bergmann’s patterns at the intraspecific (BR-I) and the community scales (CM-I and CM-A), but not when the metric did not include local trait or abundance (i.e. interspecific approach; BR-S). Interestingly, the community scale pattern which did not include local species abundances (i.e. CM-S) was contrary to that of abundance-based metrics, i.e. community species mean decreased with elevation. Some of the more abundant hawkmoth species for which we could test the intraspecific pattern individually showed a variety of patterns including positive, negative and no correlation. On the other hand, we detected the converse-Bergmann’s pattern in birds at all investigated taxonomic scales (i.e. BR-S, BR-CM-S & BR-CM-A). However, statistically it was strongest for the abundance-weighted community mean and the least for the interspecific approach. In the absence of traits of individual birds we could not test for the metrics which required individual trait measurements. Clearly, thermoregulation is not the only or even the primary mechanism influencing the body size-temperature relationship, even in endotherms. More importantly, overall, all metrics which incorporate local traits and/or abundance show stronger correlations of body mass with elevation, than when this information is lacking.

The variety in the observed patterns of the body size-temperature relationship has largely been attributed to life-history related traits of different taxa (*e*.*g*. endotherms vs ectotherms; mesotherms vs exotherms). However, our study provides evidence for differing patterns for the same taxon, across different taxonomic scales (intraspecific, interspecific or community), and with different metrics (e.g. with or without abundance; with or without individual trait measurements), even when using the same data set. Clearly, the scale and the metric with which Bergmann’s pattern is tested influences the conclusion of a study; i.e. each combination of metric and scale, presumably reflecting a different process, has its own body size-temperature pattern. Therefore, it may be best to treat ‘Bergmann’s Rule’ simply as a label for a pattern associated with the negative relationship between body size and temperature; identifying the underlying process necessarily requires an understanding of the metric and the scale involved, as also an appreciation of how the pattern changes with these two, even for the same data.

Along with taxonomic scales, another important consideration when comparing body size variation across studies is the difference in the spatial scales (Levins 1992; Chase & Knight 2013; Chase et al. 2018). Most studies of Bergmann’s pattern have been conducted across continental scales (i.e., latitudinal gradients; Olalla-Tárraga et al. 2006; Rodriíguez et al. 2008; Ollala-Tarraga et al. 2010; Zeuss et al. 2017). It has been suggested that the dominance of species turnover obviates the need to incorporate intraspecific or abundance variation at latitudinal scales, whereas intra-community dynamics, such as competition, play a stronger role in facilitating co-existence for shorter elevational transects where the environment changes substantially within a small geographic scale (Siefert et al. 2015). However, a large number of intraspecific analyses of Bergmann’s patterns have shown that many species can exhibit substantial phenotypic or genetic variation in body mass even across continental scales (Ashton 2002; Adams & Church 2007; Hassall 2015; Goldberg et al. 2018). To our best knowledge, no study has yet assessed the role of intraspecific/abundance variation in determining community-level Bergmann’s patterns across broad latitudinal gradients, although examples exist in the plant functional ecology literature (Siefert et al. 2015).

In our study, which spans a geographically compact transect, metrics which used abundance show higher significance of correlation, more so for hawkmoths than for birds. The difference in role of species abundance between hawkmoths and birds may be related to their beta diversities: 0.68 for hawkmoths and 0.93 for birds (Sørensen Indices; Supporting Information Figure S3), suggesting the presence of species with broader rangers in the former. Intraspecific variation and change in abundance may play a greater role in the change in community mean trait in taxa containing species with large geographical ranges. This also corroborates the lack of any significant correlation in the BR-S and BR-CM-S approach for hawkmoths, both of which lacked species relative abundances. On the other hand, turnover in species occurrences dominated the community-mean variation in birds and hence the negative body mass-elevation correlation was detected with and without abundance in both BR-S and BR-C.

Further, while latitudinal and elevational transects are most often viewed as similar through the prism of temperature, they differ in several important, but rarely considered, determinants of animal body size like oxygen availability (Körner 2007), air density (Dillon et al. 2006), UV-radiation (Mani 1968), etc. which may have contrasting influence on organismal body size through multiple pathways (Klepsatel et al. 2014). For instance, temperature seasonality is more prominent at high latitudes than at high elevations in the tropics (Rahbeck 1997; Körner 2000). Contrasting patterns of body size variation between latitudinal and elevational gradients has been reported for both endotherms and ectotherms (Gutiérrez-Pinto et al. 2014; Klepsatel et al. 2014; Sun et al. 2017) and therefore, coherence between latitudinal and elevational Bergmann’s patterns is not always to be expected.

### Bergmann’s patterns in hawkmoths

At the intraspecific scale, the co-added data from individuals of all hawkmoth species exhibited a weak but statistically significant positive Bergmann’s pattern. It should be noted that this is the average intraspecific value for the entire community. Of the 24 species tested separately, 6 (25%) exhibited a significant Bergmann’s pattern while 1 species exhibited a converse Bergmann’s pattern. Two of these (*Cechetra lineosa* and *Acosmeryx naga)*, represent the most abundant species in the region (> 25% in our data set) and span almost the entire sampled elevational extent. It has been previously remarked that intraspecific clines in body size are generally expected only for the ‘common’ species with large geographic ranges (*sensu* Chown & Gaston 2010). One reason for this may be the lack of statistical power for the smaller samples of the less abundant species for a given effort. Another reason for this may come from the generally smaller geographical, and hence environmental ranges associated with rare species. Therefore the intrinsic dispersion in the trait values within such a species would be a larger fraction of the width in environmental optimum across the geographical range. Whether the intraspecifc Bergmann’s clines in species of hawkmoths of our study region are due to genetic adaptation or phenotypic plasticity remains to be investigated.

At the species level, we did not find any significant correlation between body mass and elevation, which is in agreement with the three previous studies of the interspecific Bergmann’s pattern in moths: of hawkmoths across latitude (Beerli et al. 2019), of Geometridae across elevation (Brehm & Fiedler 2004), and of macrolepidoptera in general across elevation (Beck et al. 2016). Beerli et al. (2019) reported a support for the resource availability hypothesis for hawkmoths at a latitudinal scale which links adult body size to resource availability during the developmental phase (Rosenzweig 1968). Our results do not support this, since productivity was strongly negatively correlated with elevation along our study transect (NDVI ∼ elevation: *r*^*2*^ *= 0*.*84; p < 0*.*01*).

No previous work has assessed community level variation in body size for the hawkmoth family. Brehm & Fiedler (2004) found no significant trend for the family *Geometridae* at the community level, although their smaller sampled elevational range (1000-2700 m) may have made the detection more difficult. On the other hand, Zeuss et al. (2017) found a converse-Bergmann’s pattern for lepidopteran assemblages at a continental scale. They showed that voltinism, or the number of generations per year, may limit adult body size in macrolepidoptera by affecting the time available for development during the larval stage, although with species-specific exceptions. We note that the non-abundance weighted community-mean metric yielded a converse-Bergmann’s pattern in our study also, though we are not in a position to test for voltinism at this stage. Brehm et al. (2019) reported a positive Bergmann’s pattern in the ectothermic *Geometridae* and *Arctiinae* moths in Costa Rica, and ruled out thermoregulation as the primary mechanism. It has been suggested that hawkmoths should be considered as (behavioural) endotherms – for the purpose of Bergmann’s pattern – because of their active ‘shivering’ of thoracic muscles to heat up the body (Heinrich 1993; Beerli et al. 2019) which explains the positive body size-elevation relationship at the community level in our study. However, it may be noted that the only other community-approach to Bergmann’s patterns in moths, which incorporates both – intraspecific variation and species relative abundances – and was conducted along a tropical elevational gradient for a direct comparison, also revealed a strong positive Bergmann’s patterns along the elevational gradient in strictly ectothermic groups i.e. *Geometridae* and *Arctiinae* (Brehm et al. 2019). This raises an important question of whether this similarity is methodological, or mechanistic.

Overall, interspecific variation of species turnover accounted for an average 54%, where as intraspecific variation accounted for 33% of the total community-level body mass variance in hawkmoths as shown through variance partitioning. This result is consistent with a growing body of literature advocating the use of both individual and species-specific traits to investigate community level trait-environment relationships (Jung et al. 2014, Siefert et al. 2015, Enquist et al. 2015; Classen et al. 2017). Only few studies have explored the extent of intraspecific variability in insect communities till date, and obtained very contrasting values (e.g. < 5% for dung beetles (*Insecta: Coleoptera*) by Griffiths et al. 2016; < 1% for stonefly assemblages (*Insecta: Plecoptera*) by Garcia-Raventós et al. 2017; and > 70% for spider communities (Arachnida: Araneae) by Dahirel et al. 2017). Interestingly, our results are very similar to the values reported for several plant communities (reviewed in Auger & Shipley 2013 and global meta-analysis in Siefert et al. 2015). Other taxa have reported mixed results (33% for tadpoles by Xavier Jordani et al. 2019; 70% in lichens by Asplund & Wardle 2014). It is suggested that community-level trait variation, especially across broad environmental gradients, is driven primarily by species turnover, but the relative importance of intraspecific variation depends strongly on the trait, environmental factor, and spatial scale considered (Araújo & Costa-Pereira 2013; Siefert etal. 2014). In the context of Bergmann’s pattern, our results suggest that intraspecific variation should be an important consideration for groups with low beta diversity (i.e. large ranges of species within the study site), and at shorter spatial scales (e.g. local elevational transects). However, more studies will be needed to affirm the generality of this observation.

### Converse-Bergmann’s Patterns in Birds

We found a converse Bergmann’s pattern in birds at both community and interspecific scales. This is entirely against the expectation from the classical thermoregulatory mechanism in endothermic organisms, but matches with the previous (interspecific) pattern observed in the region (Schumm et al. 2020). Schumm et al. (2020) reported a decreasing trend for forest species and positive relationship for open habitat species and an initial decrease followed by increase above 3500 m elevation for the entire community. We note that this transition in pattern occurred above our sampling limit. Freeman (2017) found no significant relationship between species body mass and elevational range mid-point for passerines in New Guinea, Peru and Costa Rica and suggested that thermoregulation may not be the primary determinant for avian body mass along tropical elevational transects.

Birds are among the most studied of animal groups and ecologists have a better understanding of the various life-history traits that characterise this speciose group. Not surprisingly, Bergmann’s pattern has been explored along multiple life-history dimensions to understand their impact on the basic thermoregulatory paradigm. We have already discussed the issue of beta diversity impacting Bergmann’s pattern. While thermoregulation may cause body size to increase with decreasing temperature, resource (food) availability may result in a positive correlation due to reduced primary productivity at low temperatures. In this context, it has been reported that the decrease of avian body size with elevation in the eastern Himalayas is correlated with decreasing arthropod (their prey) body size (Price et al. 2014; Schumm et al. 2020). But, results from Brehm et al. (2019) and this work show that all insect groups do not show such a decrease, especially when local abundances are incorporated.

The expected Bergmann’s pattern has been seen more regularly in birds across latitudinal gradients, but even here the results suggest the modifying influence of life-history traits: weaker pattern for species which are migratory and/or have sheltered nesting in the western Palearctic (Mainwaring & Street 2019); contrasting pattern for endemics and non-endemics in the Andes, both along latitudinal and elevational gradients; successful establishment of alien species is less dependent on thermoregulatory considerations than the location of introduction (Blackburn et al. 2019). On the other hand, Olson et al. (2009) concluded from a global analysis of over 9000 species that the positive Bergmann’s pattern was driven by a combination of temperature and resource availability.

Analysing a large data set of North American birds, Goodman et al. (2012) found that body size was more impacted by climatic variability and primary productivity than mean ambient temperature. We re-emphasize that taxonomic and spatial scales are important considerations for comparing mechanistic processes, however we did not find any previous study that has assessed body size variation in birds along elevational transects using abundance-weighted community mean, for a direct comparison.

## Conclusion

Despite a large body of literature on characterizing Bergmann’s patterns, no study has yet empirically demonstrated geographic gradients in body size at the population, species and community level using the same data. Our cross-taxon and cross-scale comparison using hawkmoths and birds along the same 2600 m elevational gradient revealed discrepant patterns for the two taxa, and at different taxonomic scales. We suggest that drawing taxon-specific generalization of processes, on the basis of results from across taxonomic scales is premature. We further highlight the importance of species abundances and intraspecific variation for characterizing geographic gradients in body size at the level of communities, and suggest that the validity of the current approach of using assemblage-means based on species occurrences and mean body sizes needs to be empirically verified, especially along elevational gradients that may span the entire geographic ranges of several species within a taxa. Methodologically, we argue for a community-oriented approach in future studies for addressing the generality of Bergmann’s pattern because of its potential in resolving the relative importance of individual and species-level processes. Our study, though not conducted to explicitly test any particular mechanism, provides strong support against the classical thermoregulatory mechanism in endotherms, and warrants further investigations into the likely mechanism (including heat conservation) governing body size patterns for ectotherms. More in-depth analysis across different taxonomic scales, especially in a comparative frame-work across different taxa along the same transect are required for an accurate charachterization of the Bergmann’s pattern. We conclude by reiterating the key lesson from this study: that every study should consider, and address, the taxonomic- and spatial-scale dependency of Bergmann’s patterns, and species relative abundance and intraspecific trait variation may be essential ingredients for doing so.

## Notes

### Competing Interest Statement

The authors have declared no competing interest.

## References

1. Adams, D.C. and Church, J.O., 2008. Amphibians do not follow Bergmann’s rule. Evolution: International Journal of Organic Evolution, 62(2), pp.413–420.

2. Alhajeri, B.H. and Steppan, S.J., 2016. Association between climate and body size in rodents: a phylogenetic test of Bergmann’s rule. Mammalian Biology, 81(2), pp.219–225.

3. Ali, S., and S. D. Ripley. 1972. Handbook of the birds of India and Pakistan, together with those of Bangladesh, Nepal, Bhutan, and Sri Lanka. Oxford University Press, Bombay.

4. Araújo, M.S. and Costa-Pereira, R., 2013. Latitudinal gradients in intraspecific ecological diversity. Biology letters, 9(6), p.20130778.

5. Ashton, K.G., 2002. Patterns of within-species body size variation of birds: strong evidence for Bergmann’s rule. Global Ecology and Biogeography, 11(6), pp.505–523.

6. Asplund, J. and Wardle, D.A., 2014. Within species variability is the main driver of community level responses of traits of epiphytes across a long term chronosequence. Functional Ecology, 28(6), pp.1513–1522.

7. Atkinson, D. and Sibly, R.M., 1997. Why are organisms usually bigger in colder environments? Making sense of a life history puzzle. Trends in ecology & evolution, 12(6), pp.235–239.

8. Auger, S. and Shipley, B., 2013. Inter specific and intra specific trait variation along short environmental gradients in an old growth temperate forest. Journal of Vegetation Science, 24(3), pp.419–428.

9. Bartholemew, G.A., Vleck, D. and Vleck, C.M., 1981. Instantaneous measurements of oxygen consumption during pre-flight warm-up and post-flight cooling in sphingid and saturnid moths. Journal of Experimental Biology, 90, pp.17–32.

10. Beck, J. and Kitching, I.J., 2009. Drivers of moth species richness on tropical altitudinal gradients: a cross-regional comparison. Global Ecology and Biogeography, 18(3), pp.361–371.

11. Beck, J., Kitching, I.J. and Linsenmair, K.E., 2006a. Wallace’s line revisited: has vicariance or dispersal shaped the distribution of Malesian hawkmoths (Lepidoptera: Sphingidae)?. Biological Journal of the Linnean Society, 89(3), pp.455–468.

12. Beck, J., Kitching, I.J. and Linsenmair, K.E., 2006b. Effects of habitat disturbance can be subtle yet significant: biodiversity of hawkmoth-assemblages (Lepidoptera: Sphingidae) in Southeast-Asia. In Arthropod Diversity and Conservation (pp. 451–472). Springer, Dordrecht.

13. Beck, J., Liedtke, H.C., Widler, S., Altermatt, F., Loader, S.P., Hagmann, R., Lang, S. and Fiedler, K., 2016. Patterns or mechanisms? Bergmann’s and Rapoport’s rule in moths along an elevational gradient. Community Ecology, 17(2), pp.137–148.

14. Beerli, N., Bärtschi, F., Ballesteros-Mejia, L., Kitching, I.J. and Beck, J., 2019. How has the environment shaped geographical patterns of insect body sizes? A test of hypotheses using sphingid moths. Journal of Biogeography, 46(8), pp.1687–1698.

15. Bidau, C.J. and Martí, D.A., 2007. Clinal variation of body size in Dichroplus pratensis (Orthoptera: Acrididae): inversion of Bergmann’s and Rensch’s rules. Annals of the Entomological Society of America, 100(6), pp.850–860.

16. Bivand, Roger S. and Wong, David W. S. (2018) Comparing implementations of global and local indicators of spatial association TEST, 27(3), 716-748. URL https://doi.org/10.1007/s11749-018-0599-x

17. Blackburn, T.M. and Gaston, K.J., 1996. Spatial patterns in the body sizes of bird species in the New World. Oikos, pp.436–446.

18. Blackburn, T.M. and Hawkins, B.A., 2004. Bergmann’s rule and the mammal fauna of northern North America. Ecography, 27(6), pp.715–724.

19. Blackburn, T.M., Redding, D.W. and Dyer, E.E., 2019. Bergmann’s rule in alien birds. Ecography, 42(1), pp.102–110.

20. Brehm, G. and Fiedler, K., 2004. Bergmann’s rule does not apply to geometrid moths along an elevational gradient in an Andean montane rain forest. Global Ecology and Biogeography, 13(1), pp.7–14.

21. Brehm, G., Zeuss, D. and Colwell, R.K., 2019. Moth body size increases with elevation along a complete tropical elevational gradient for two hyperdiverse clades. Ecography, 42(4), pp.632–642.

22. Chase, J.M. and Knight, T.M., 2013. Scale dependent effect sizes of ecological drivers on biodiversity: why standardised sampling is not enough. Ecology letters, 16, pp.17–26.

23. Chase, J.M., McGill, B.J., McGlinn, D.J., May, F., Blowes, S.A., Xiao, X., Knight, T.M., Purschke, O. and Gotelli, N.J., 2018. Embracing scale dependence to achieve a deeper understanding of biodiversity and its change across communities. Ecology letters, 21(11), pp.1737–1751.

24. Chown, S.L. and Gaston, K.J., 2010. Body size variation in insects: a macroecological perspective. Biological Reviews, 85(1), pp.139–169.

25. Classen, A., Steffan-Dewenter, I., Kindeketa, W.J. and Peters, M.K., 2017. Integrating intraspecific variation in community ecology unifies theories on body size shifts along climatic gradients. Functional Ecology, 31(3), pp.768–777.

26. Cornwell, W.K. and Ackerly, D.D., 2009. Community assembly and shifts in plant trait distributions across an environmental gradient in coastal California. Ecological Monographs, 79(1), pp.109–126.

27. Cushman, J.H., Lawton, J.H. and Manly, B.F., 1993. Latitudinal patterns in European ant assemblages: variation in species richness and body size. Oecologia, 95(1), pp.30–37

28. Dahirel, M., Dierick, J., De Cock, M. and Bonte, D., 2017. Intraspecific variation shapes community level behavioral responses to urbanization in spiders. Ecology, 98(9), pp.2379–2390.

29. de la Riva, E.G., Pérez Ramos, I.M., Tosto, A., Navarro Fernández, C.M., Olmo, M., Marañón, T. and Villar, R., 2016. Disentangling the relative importance of species occurrence, abundance and intraspecific variability in community assembly: a trait based approach at the whole plant level in Mediterranean forests. Oikos, 125(3), pp.354–363.

30. Des Roches, S., Post, D.M., Turley, N.E., Bailey, J.K., Hendry, A.P., Kinnison, M.T., Schweitzer, J.A. and Palkovacs, E.P., 2018. The ecological importance of intraspecific variation. Nature ecology & evolution, 2(1), pp.57–64.

31. Dillon, M.E., Frazier, M.R. and Dudley, R., 2006. Into thin air: physiology and evolution of alpine insects. Integrative and Comparative Biology, 46(1), pp.49–61.

32. Diniz-Filho, J.A.F., Bini, L.M. and Hawkins, B.A., 2003. Spatial autocorrelation and red herrings in geographical ecology. Global ecology and Biogeography, 12(1), pp.53–64.

33. Enquist, B.J., Norberg, J., Bonser, S.P., Violle, C., Webb, C.T., Henderson, A., Sloat, L.L. and Savage, V.M., 2015. Scaling from traits to ecosystems: developing a general trait driver theory via integrating trait-based and metabolic scaling theories. Advances in ecological research, 52, pp.249–318.

34. Fan, L., Cai, T., Xiong, Y., Song, G. and Lei, F., 2019. Bergmann’s rule and Allen’s rule in two passerine birds in China. Avian Research, 10(1), pp.1–11.

35. Freeman, B.G., 2017. Little evidence for Bergmann’s rule body size clines in passerines along tropical elevational gradients. Journal of Biogeography, 44(3), pp.502–510.

36. Garcia Raventós, A., Viza, A., Tierno de Figueroa, J.M., Riera, J.L. and Múrria, C., 2017. Seasonality, species richness and poor dispersion mediate intraspecific trait variability in stonefly community responses along an elevational gradient. Freshwater Biology, 62(5), pp.916–928.

37. Gaston, K.J., Chown, S.L. and Evans, K.L., 2008. Ecogeographical rules: elements of a synthesis. Journal of Biogeography, 35(3), pp.483–500.

38. Gillooly, J.F., Allen, A.P., West, G.B. and Brown, J.H., 2005. The rate of DNA evolution: effects of body size and temperature on the molecular clock. Proceedings of the National Academy of Sciences, 102(1), pp.140–145.

39. Gohli, J. and Voje, K.L., 2016. An interspecific assessment of Bergmann’s rule in 22 mammalian families. BMC Evolutionary Biology, 16(1), pp.1–12.

40. Goldberg, J., Cardozo, D., Brusquetti, F., Bueno Villafañe, D., Caballero Gini, A. and Bianchi, C., 2018. Body size variation and sexual size dimorphism across climatic gradients in the widespread treefrog Scinax fuscovarius (Anura, Hylidae). Austral Ecology, 43(1), pp.35–45.

41. Goodman, R.E., Lebuhn, G., Seavy, N.E., Gardali, T. and Bluso-Demers, J.D., 2012. Avian body size changes and climate change: warming or increasing variability?. Global Change Biology, 18(1), pp.63–73.

42. Griffiths, H.M., Louzada, J., Bardgett, R.D. and Barlow, J., 2016. Assessing the importance of intraspecific variability in dung beetle functional traits. PloS one, 11(3), p.e0145598.

43. Gutiérrez-Pinto, N., McCracken, K.G., Alza, L., Tubaro, P., Kopuchian, C., Astie, A. and Cadena, C.D., 2014. The validity of ecogeographical rules is context-dependent: testing for Bergmann’s and Allen’s rules by latitude and elevation in a widespread Andean duck. Biological Journal of the Linnean Society, 111(4), pp.850–862.

44. Hassall, C., 2015. Odonata as candidate macroecological barometers for global climate change. Freshwater Science, 34(3), pp.1040–1049.

45. Heinrich, 8., 1993: The Hot-Blooded Insects. Strategies and Mechanisms of Thermoregulation. Berlin and Heidelberg: Springer-Verlag. 60 I pp.

46. Oksanen J., Blanchet G. F., Friendly M., Kindt R., Legendre P., McGlinn D., Minchin P. R., R. B. O’Hara, Simpson G. L., Solymos P., M. Henry H. Stevens, Szoecs E. and Wagner H. (2019). vegan: Community Ecology Package. R package version 2.5-6. https://CRAN.R-project.org/package=vegan

47. Jetz, W., Thomas, G.H., Joy, J.B., Hartmann, K. and Mooers, A.O., 2012. The global diversity of birds in space and time. Nature, 491(7424), pp.444–448.

48. Jung, V., Albert, C.H., Violle, C., Kunstler, G., Loucougaray, G. and Spiegelberger, T., 2014. Intraspecific trait variability mediates the response of subalpine grassland communities to extreme drought events. Journal of ecology, 102(1), pp.45–53.

49. Keddy, P.A., 1992. Assembly and response rules: two goals for predictive community ecology. Journal of vegetation science, 3(2), pp.157–164.

50. Kichenin, E., Wardle, D.A., Peltzer, D.A., Morse, C.W. and Freschet, G.T., 2013. Contrasting effects of plant inter-and intraspecific variation on community-level trait measures along an environmental gradient. Functional Ecology, 27(5), pp.1254–1261.

51. Kitching, I.J. 2020. Sphingidae Taxonomic Inventory, http://sphingidae.myspecies.info/, accessed on 15 November 2019

52. Klepsatel, P., Gáliková, M., Huber, C.D. and Flatt, T., 2014. Similarities and differences in altitudinal versus latitudinal variation for morphological traits in Drosophila melanogaster. Evolution, 68(5), pp.1385–1398.

53. Körner, C. 2000. Why are there global gradients in species richness? Mountains might hold the answer. Trends Ecol. Evol. 15: 513514.

54. Kuhn M., 2020. caret: Classification and Regression Training. R package version 6.0-86. https://CRAN.R-project.org/package=caret

55. Kuznetsova A, Brockhoff PB, Christensen RHB (2017). “lmerTest Package: Tests in Linear Mixed Effects Models.” _Journal of Statistical Software_, *82*(13), 1–26. doi: 10.18637/jss.v082.i13 (URL: https://doi.org/10.18637/jss.v082.i13).

56. Lepš, J., de Bello, F., Šmilauer, P. and Doležal, J., 2011. Community trait response to environment: disentangling species turnover vs intraspecific trait variability effects. Ecography, 34(5), pp.856–863.

57. Levin, S.A., 1992. The problem of pattern and scale in ecology: the Robert H. MacArthur award lecture. Ecology, 73(6), pp.1943–1967.

58. MacArthur, R. and Levins, R., 1967. The limiting similarity, convergence, and divergence of coexisting species. The american naturalist, 101(921), pp.377–385.

59. Mainwaring, M.C. and Street, S.E., 2019. Conformity to Bergmann’s rule in birds depends on nest design and migration. bioRxiv, p.686972.

60. Mani, M.S., 1968. Ecological specializations of high altitude insects. In Ecology and biogeography of high altitude insects (pp. 51–74). Springer, Dordrecht.

61. McNab, B.K., 2010. Geographic and temporal correlations of mammalian size reconsidered: a resource rule. Oecologia, 164(1), pp.13–23.

62. Meiri, S. and Dayan, T., 2003. On the validity of Bergmann’s rule. Journal of biogeography, 30(3), pp.331–351.

63. Meiri, S. and Thomas, G.H., 2007. The geography of body size–challenges of the interspecific approach. Global Ecology and Biogeography, 16(6), pp.689–693.

64. Mousseau, T.A., 1997. Ectotherms follow the converse to Bergmann’s rule. Evolution, pp.630–632.

65. Mungee, M. and Athreya, R., 2020a. Rapid photogrammetry of morphological traits of free-ranging moths. Ecological Entomology, 45(5), pp.911–923.

66. Mungee, M. and Athreya, R., 2020b. Intraspecific trait variability and community assembly in hawkmoths (Lepidoptera: Sphingidae) across an elevational gradient in the eastern Himalayas, India. Ecology & Evolution; 10.1002/ece3.7054

67. Myers, N., Mittermeier, R.A., Mittermeier, C.G., Da Fonseca, G.A. and Kent, J., 2000. Biodiversity hotspots for conservation priorities. Nature, 403(6772), pp.853–858.

68. Nwaogu, C.J., Tieleman, B.I., Bitrus, K. and Cresswell, W., 2018. Temperature and aridity determine body size conformity to Bergmann’s rule independent of latitudinal differences in a tropical environment. Journal of ornithology, 159(4), pp.1053–1062.

69. Olalla-Tárraga, M.Á. and Rodríguez, M.Á., 2007. Energy and interspecific body size patterns of amphibian faunas in Europe and North America: anurans follow Bergmann’s rule, urodeles its converse. Global Ecology and Biogeography, 16(5), pp.606–617.

70. Olalla-Tárraga, M.Á., 2011. “Nullius in Bergmann” or the pluralistic approach to ecogeographical rules: a reply to Watt et al.(2010). Oikos, 120(10), pp.1441–1444.

71. Olalla-Tárraga, M.A., Bini, L.M., Diniz-Filho, J.A. and Rodríguez, M.Á., 2010. Cross-species and assemblage-based approaches to Bergmann’s rule and the biogeography of body size in Plethodon salamanders of eastern North America. Ecography, 33(2), pp.362–368.

72. Olalla-Tárraga, M.Á., Diniz-Filho, J.A.F., Bastos, R.P. and Rodríguez, M.Á., 2009. Geographic body size gradients in tropical regions: water deficit and anuran body size in the Brazilian Cerrado. Ecography, 32(4), pp.581–590

73. Olalla-Tárraga, M.Á., Rodríguez, M.Á. and Hawkins, B.A., 2006. Broad-scale patterns of body size in squamate reptiles of Europe and North America. Journal of Biogeography, 33(5), pp.781–793.

74. Olson, V.A., Davies, R.G., Orme, C.D.L., Thomas, G.H., Meiri, S., Blackburn, T.M., Gaston, K.J., Owens, I.P. and Bennett, P.M., 2009. Global biogeography and ecology of body size in birds. Ecology Letters, 12(3), pp.249–259.

75. Orme, C.D.L., Davies, R.G., Burgess, M., Eigenbrod, F., Pickup, N., Olson, V.A., Webster, A.J., Ding, T.S., Rasmussen, P.C., Ridgely, R.S. and Stattersfield, A.J., 2005. Global hotspots of species richness are not congruent with endemism or threat. Nature, 436(7053), pp.1016–1019.

76. Pagel M. Inferring the historical patterns of biological evolution. Nature. 1999;401:877–84.

77. Paradis E. & Schliep K. 2018., ape 5.0: an environment for modern phylogenetics and evolutionary analyses in R. Bioinformatics 35: 526–528.

78. Price, T.D., 2012. Eaglenest Wildlife Sanctuary: Pressures on Biodiversity: (EO Wilson Award Address).

79. Price, T.D., Hooper, D.M., Buchanan, C.D., Johansson, U.S., Tietze, D.T., Alström, P., Olsson, U., Ghosh-Harihar, M., Ishtiaq, F., Gupta, S.K. and Martens, J., 2014. Niche filling slows the diversification of Himalayan songbirds. Nature, 509(7499), pp.222–225.

80. Quintero, I. and Jetz, W., 2018. Global elevational diversity and diversification of birds. Nature, 555(7695), pp.246–250.

81. Rahbek, C., 1997. The relationship among area, elevation, and regional species richness in neotropical birds. The American Naturalist, 149(5), pp.875–902.

82. Rasmussen, P. C., and J. C. Anderton. 2005. Birds of south Asia: The Ripley guide, Vol. 2. Lynx edicions, Barcelona, Spain.

83. Riemer, K., Guralnick, R.P. and White, E.P., 2018. No general relationship between mass and temperature in endothermic species. Elife, 7, p.e27166.

84. Rodríguez, M.Á., Olalla-Tárraga, M.Á. and Hawkins, B.A., 2008. Bergmann’s rule and the geography of mammal body size in the Western Hemisphere. Global Ecology and Biogeography, 17(2), pp.274–283.

85. Rosenzweig, M.L., 1968. Net primary productivity of terrestrial communities: prediction from climatological data. The American Naturalist, 102(923), pp.67–74.

86. Salewski, V. and Watt, C., 2017. Bergmann’s rule: a biophysiological rule examined in birds. Oikos, 126(2).

87. Schumm, M., White, A.E., Supriya, K. and Price, T.D., 2020. Ecological limits as the driver of bird species richness patterns along the east Himalayan elevational gradient. The American Naturalist, 195(5), pp.802–817.

88. Shelomi, M., 2012. Where are we now? Bergmann’s rule sensu lato in insects. The American Naturalist, 180(4), pp.511–519.

89. Siefert, A., Violle, C., Chalmandrier, L., Albert, C.H., Taudiere, A., Fajardo, A., Aarssen, L.W., Baraloto, C., Carlucci, M.B., Cianciaruso, M.V. and de L. Dantas, V., 2015. A global meta-analysis of the relative extent of intraspecific trait variation in plant communities. Ecology letters, 18(12), pp.1406–1419.

90. Sun, Y., Li, M., Song, G., Lei, F., Li, D. and Wu, Y., 2017. The role of climate factors in geographic variation in body mass and wing length in a passerine bird. Avian Research, 8(1), pp.1–9.

91. Taudiere, A. and Violle, C., 2016. cati: an R package using functional traits to detect and quantify multi-level community assembly processes. Ecography, 39(7), pp.699–708.

92. Vellend, M., Lajoie, G., Bourret, A., Múrria, C., Kembel, S.W. and Garant, D., 2014. Drawing ecological inferences from coincident patterns of population-and community-level biodiversity. Molecular Ecology, 23(12), pp.2890–2901.

93. Watt, C., Mitchell, S. and Salewski, V., 2010. Bergmann’s rule; a concept cluster?. Oikos, 119(1), pp.89–100.

94. Wilman, H., Belmaker, J., Simpson, J., de la Rosa, C., Rivadeneira, M.M. and Jetz, W., 2014. EltonTraits 1.0: Species-level foraging attributes of the world’s birds and mammals: Ecological Archives E095-178. Ecology, 95(7), pp.2027–2027.

95. Xavier Jordani, M., Mouquet, N., Casatti, L., Menin, M., de Cerqueira Rossa Feres D. and Albert, C.H., 2019. Intraspecific and interspecific trait variability in tadpole meta communities from the Brazilian Atlantic rainforest. Ecology and evolution, 9(7), pp.4025–4037.

96. Yom-Tov, Y. and Geffen, E., 2006. Geographic variation in body size: the effects of ambient temperature and precipitation. Oecologia, 148(2), pp.213–218.

97. Zamora Camacho, F.J., Reguera, S. and Moreno Rueda, G., 2014. Bergmann’s Rule rules body size in an ectotherm: heat conservation in a lizard along a 2200 metre elevational gradient. Journal of Evolutionary Biology, 27(12), pp.2820–2828.

98. Zeuss, D., Brunzel, S. and Brandl, R., 2017. Environmental drivers of voltinism and body size in insect assemblages across Europe. Global Ecology and Biogeography, 26(2), pp.154–165.

99. Zug, G.R., Vitt, L.J. & Caldwell, J.P. (2001) Herpetology. Academic Press, San Diego

