## Supplementary_01 for "Taxonomic scale dependency of Bergmann’s patterns: A cross-scale comparison of hawkmoths and birds along a tropical elevational gradient"

### Online Supporting Information
#### Appendix S1

**Table S1.** Summary statistics for hawkmoth and bird data. N = number of individuals recorded, $S_{obs}$ = Number of species observed, $S_{rare}$ = Rarefied species richness; Alpha = Species diversity calculated using Fisher's alpha diversity index

| Elevation | N | | $S_{obs}$ | | $S_{rare}$ | | Alpha | |
|---|---|---|---|---|---|---|---|---|
| | Hawkmohts | Birds | Hawkmohts | Birds | Hawkmohts | Birds | Hawkmohts | Birds |
| **200** | 473 | 549 | 43 | 53 | 39 | 53 | 11.5 | 15 |
| **500** | 378 | 820 | 39 | 81 | 37 | 74 | 10.9 | 22 |
| **700** | 335 | 977 | 33 | 84 | 33 | 76 | 09.1 | 22 |
| **900** | 347 | 852 | 45 | 77 | 44 | 70 | 13.8 | 21 |
| **1100** | 350 | 1095 | 48 | 88 | 46 | 76 | 15.1 | 23 |
| **1300** | 332 | 1276 | 31 | 82 | 31 | 73 | 08.4 | 20 |
| **1500** | 340 | 1162 | 40 | 87 | 39 | 78 | 11.8 | 22 |
| **1700** | 359 | 1264 | 36 | 85 | 35 | 74 | 10.0 | 21 |
| **1900** | 376 | 1120 | 40 | 77 | 38 | 68 | 11.3 | 19 |
| **2100** | 323 | 1543 | 29 | 73 | 29 | 63 | 07.7 | 16 |
| **2300** | 359 | 1835 | 27 | 81 | 26 | 66 | 06.8 | 17 |
| **2500** | 363 | 1755 | 23 | 77 | 22 | 62 | 05.5 | 17 |
| **2700** | 396 | 1498 | 32 | 69 | 30 | 56 | 08.2 | 15 |
| **Total** | **4,731** | **15,746** | **80** | **245** | ---- | ---- | ---- | --- |

**Table S2.** Linear regression between intraspecific body size and elevation.. Species with more than 20 individuals, and spanning at least 800 m in elevation were analysed. *N* = total number of individuals of the species; *Range (m)* = Minimum and maximum elevation at which the species was recorded; All species with slopes significant at p < 0.1 are highlighted in bold.

| | Family | Species | Slope ± SE (g m$^{-}$1) | N | Range (m) | Rsq |
|---|---|---|---|---|---|---|
| 1 | **Macroglossinae** | ***Acosmeryx naga*** | **(2.443 ± 0.556) x 10-4** | **259** | **0500 – 2770** | **0.066** |
| 2 | **Macroglossinae** | ***Acosmeryx omissa*** | **(4.168 ± 0.050) x 10-4** | **147** | **0500 – 1700** | **0.111** |
| 3 | Macroglossinae | *Acosmeryx sericeus* | (-1.105 ± 1.223) x 10-4 | 94 | 0200 – 2770 | -0.002 |
| 4 | Macroglossinae | *Acosmeryx shervillii* | (-8.406 ± 0.102) x 10-5 | 175 | 0200 – 2770 | -0.002 |
| 5 | Macroglossinae | *Ampelophaga khasiana* | (-1.661 ± 9.862) x 10-5 | 99 | 1100 – 2770 | -0.010 |
| 6 | **Macroglossinae** | ***Cechetra lineosa*** | **(1.002 ± 0.273) x 10-4** | **523** | **0200 – 2770** | **0.023** |
| 7 | Macroglossinae | *Cechetra scotti* | **(1.173 ± 0.833) x 10-4** | 125 | 0900 – 2770 | 0.008 |
| 8 | Macroglossinae | *Cechetra subangustata* | **(1.102 ± 0.879) x 10-5** | 179 | 0500 – 2770 | 0.003 |
| 9 | **Macroglossinae** | ***Eupanacra sinuata*** | (9.635 ± 4.224) x 10-5 | **112** | **1100 – 2770** | **0.036** |
| 10 | Macroglossinae | *Rhagastis castor* | (8.550 ± 8.298) x 10-5 | 39 | 0200 – 2770 | 0.002 |
| 11 | Macroglossinae | *Rhagastis confusa* | (-1.001 ± 0.669) x 10-5 | 64 | 0200 – 2770 | 0.019 |
| 12 | Macroglossinae | *Rhagastis gloriosa* | (-1.005 ± 0.854) x 10-4 | 54 | 1300 – 2770 | 0.007 |
| 13 | **Macroglossinae** | ***Rhagastis lunata*** | **(2.260 ± 0.958) x 10-4** | **88** | **1100 – 2770** | **0.050** |
| 14 | Macroglossinae | *Rhagastis olivacea* | (3.151 ± 6.937) x 10-5 | 39 | 0200 – 2300 | -0.021 |
| 15 | Macroglossinae | *Theretra boisduvalii* | (-2.699 ± 6.132) x 10-5 | 174 | 0200 – 2770 | -0.005 |
| 16 | **Macroglossinae** | ***Theretra clotho*** | **(-2.241 ± 0.641) x 10-4** | **140** | **0200 – 2770** | **0.075** |
| 17 | Macroglossinae | *Theretra nessus* | (1.253 ± 1.014) x 10-4 | 33 | 0200 – 2770 | 0.016 |
| 18 | Smerinthinae | *Ambulyx ochracea* | (2.906 ± 0.571) x 10-6 | 73 | 0200 – 2770 | -0.014 |
| 19 | Smerinthinae | *Ambulyx substrigilis* | (-7.297 ± 19.38) x 10-5 | 28 | 0200 – 1500 | -0.033 |
| 20 | Smerinthinae | *Ambulyx tobii* | (1.960 ± 1.071) x 10-5 | 21 | 0200 – 2770 | 0.105 |
| 21 | **Smerinthinae** | ***Callambulyx rubricosa*** | **(9.481 ± 5.223) x 10-4** | **26** | **0200 – 1300** | **0.084** |
| 22 | Sphinginae | *Apocalypsis velox* | (7.105 ± 6.121) x 10-4 | 29 | 0900 – 2300 | 0.012 |
| 23 | Sphinginae | *Meganoton analis* | (3.756 ± 3.385) x 10-4 | 47 | 1100 – 2770 | 0.005 |
| 24 | Sphinginae | *Psilogramma menephron* | (-1.096 ± 1.944) x 10-5 | 41 | 0200 – 2770 | -0.017 |

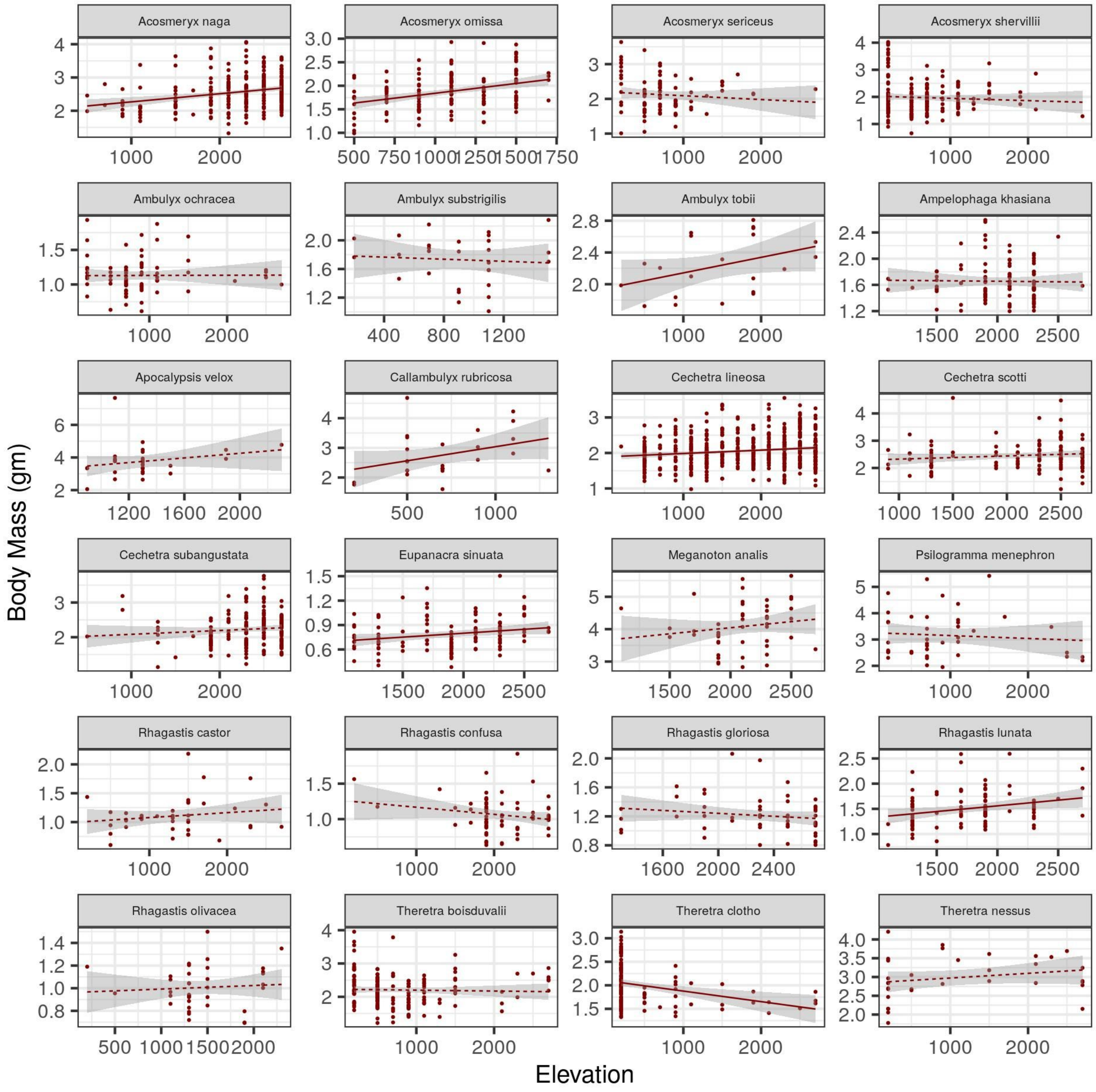


**Figure S1**. Intraspecific variation of body mass with elevation for hawkmoth species. 24 species of with more than 20 individuals and spanning more than 800 m are shown. 6 species (*Acosmeryx naga*, *Acosmeryx omissa*, *Callambulyx rubricosa*, *Cechetra lineosa*, *Eupanacra sinuata and Rhagastis lunata*) exhibited a significant intraspecific Bergmann's pattern, whereas 1 (*Theretra clotho*) exhibited a significant converse-Bergmann's clines. Regression coefficients for all the linear fits are listed in Table 3. All fits significant at $p < 0.1$ are shown as solid lines while the others are with a dashed line.

**Table S3.** Linear regression between community mean body size and elevation. The community values were weighted by rarefied species richness. All slopes were significant at $p < 0.1$.

| HAWKMOTHS | | |
|---|---|---|
| **Model** | **Slope ± SE** | **$R^2$** |
| CM-A ~ Elevation; Weighted OLS (Rarefied species richness) | **$(0.102 \pm 4.587) \times 10^{-5}$** | **0.25**** |
| CM-I ~ Elevation; Weighted OLS (Rarefied species richness) | **$(0.118 \pm 6.210) \times 10^{-5}$** | **0.18**** |
| CM-S ~ Elevation; Weighted OLS (Rarefied species richness) | **$(-0.116 \pm 5.914) \times 10^{-5}$** | **0.19*** |
| **BIRDS** | | |
| **Model** | **Slope ± SE** | **$R^2$** |
| CM-A ~ Elevation; Weighted OLS (Rarefied species richness) | **$(-3.867 \pm 0.547) \times 10^{-2}$** | **0.80***** |
| CM-S ~ Elevation; Weighted OLS (Rarefied species richness) | **$(-2.615 \pm 0.387) \times 10^{-2}$** | **0.79***** |

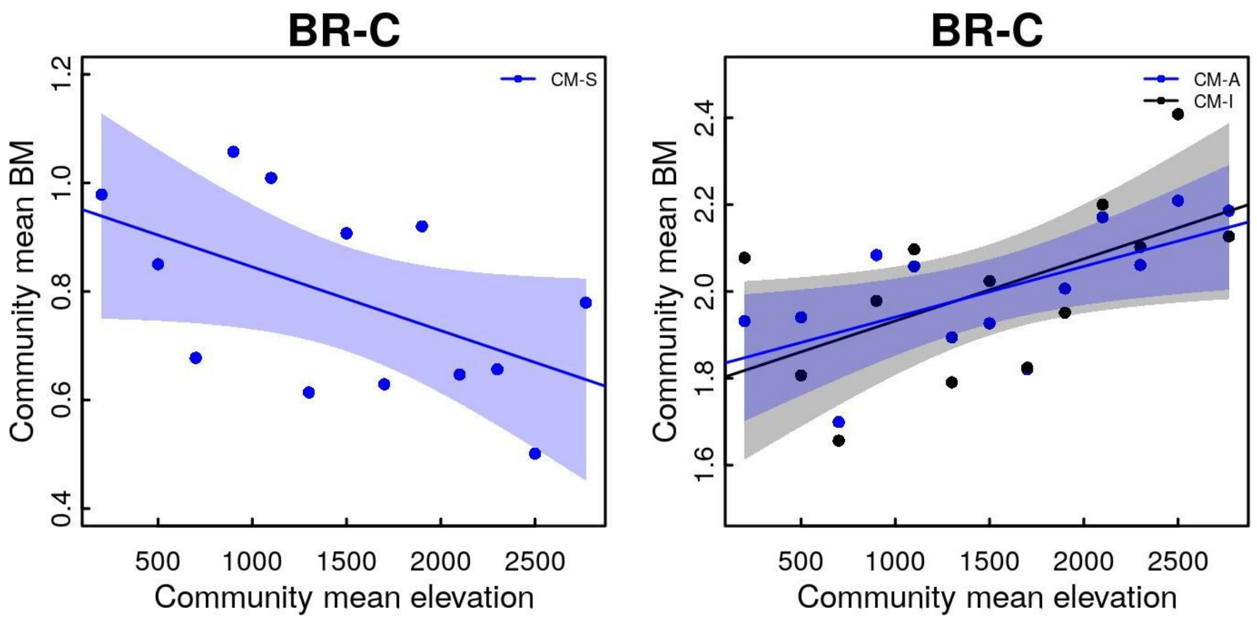


| HAWKMOTHS | | | | |
|---|---|---|---|---|
| **Taxonomic level** | **Model** | **Figure** | **Slope ± SE** | **$R^2$** |
| **Community** | CM-A ~ Elevation | S5b | **(0.117 ± 4.398) x $10^{-5}$** | **0.34**** |
| | CM-I ~ Elevation | S5b | **(0.143 ± 6.193) x $10^{-5}$** | **0.27**** |
| | CM-S ~ Elevation | S5a | **(-0.172 ± 5.692) x $10^{-5}$** | **0.21*** |

**Figure S2.** Relationship between elevation and body mass (BM) for hawkmoths at community level using the Diversity Data set; **to be compared with Main-Figure 2.** We could measure traits only for a subset (N = 3,302; 70%) of hawkmoth individuals of the complete ‘diversity data’ (N = 4,731); the remaining either did not sit on the gridded screen or were of a poor image quality. In the main manuscript, we have presented the results from the “trait data set” of 3,302 individuals. The above plots use the more complete relative abundance information from the larger “diversity data set”. We filled in this ‘missing’ trait data by randomly resampling trait values from individuals of the same species from the same elevation. **a) BR-C** (CM-S): average of the regional mean body mass of species within a community, i.e. not weighted by local abundance, **b)** BR-C (CM-A): local abundance weighted average of the regional mean body mass of species within a community; (CM-I): mean trait value of all individuals in a community). Regression coefficients for all the linear fits are listed in Table S5. All fits significant at $p < 0.1$ are shown as solid lines, the rest are as dashed lines. There are no significant differences between results from Traits and Diversity data sets (compare Main-Table 1 and Table S7) .

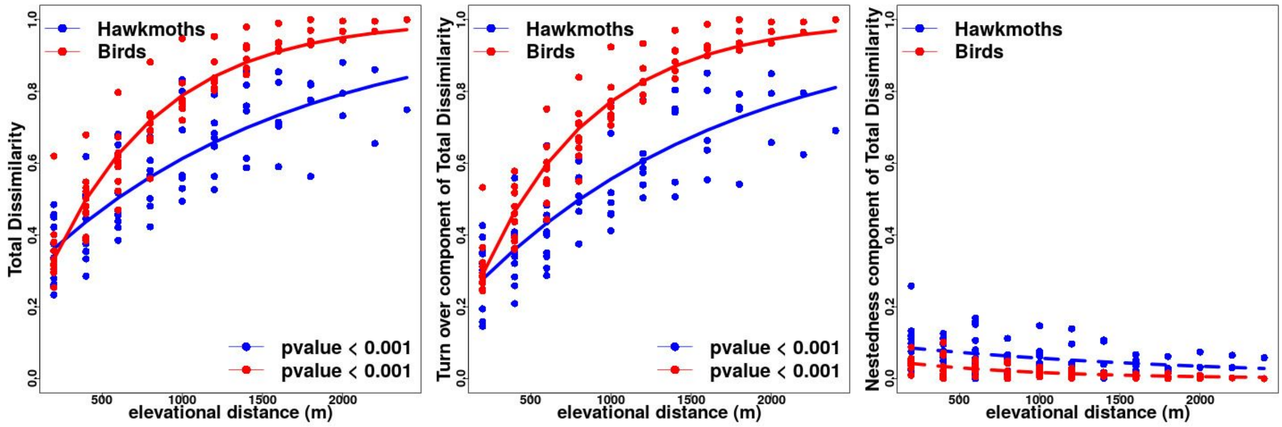


**Figure S3.** A comparison of slopes between the distance decay relationships for hawkmoths (blue) and birds (red). Sørensen index of Dissimilarity based only on presence-absence data was used to compare turnover in species occurrences, without the influence of gradients in abundance. Plots are presented for a) Total Dissimilarity, b) Turnover component of Total Dissimilarity & c). Nestedness component of Total Dissimilarity (Baselga 2010). The slope was significantly higher for birds as compared to hawkmoths (Fisher's z = 8.68, p.value < 0.001).
